# CHACAM: a cell–cell interaction–guided hierarchical attention model for high-precision cell identity annotation of scRNA-seq data in early *C. elegans* embryogenesis

**DOI:** 10.64898/2026.09.24.754260

**Authors:** Xingyu Chen, Xinxin Ju, Maithri Murali, Hongjun Li, Min Chen, Michael Q. Zhang

## Abstract

Accurate cell identity annotation is essential for reconstructing cell lineages and gene-regulatory networks in embryogenesis. However, in *Caenorhabditis elegans* (*C. elegans*) single-cell RNA-sequencing (scRNA-seq) atlases, closely related sub-lineage cells often have near-identical transcriptomes, leading to merged labels and limiting downstream analyses. We present CHACAM (Cell–Cell Interaction–guided Hierarchical Attention-based Cell Allocation Model), a supervised machine learning framework that integrates gene expression, a curated *C. elegans* ligand–receptor (L–R) interaction database, and a high-resolution cell contact map to refine cell annotations. CHACAM learns interpretable “interaction signatures” that distinguish contacting vs. non-contacting neighbors and applies a triangulation strategy using reference cells to resolve ambiguous sister identities. Applied to the Tintori and Cole–Yanai embryo atlases, CHACAM achieves near-perfect annotation accuracy, resolving longstanding ambiguities of the AB lineage and other lineages. The model also identifies new discriminative marker genes and provides mechanistic hypotheses linking contact-specific signaling to fate specification.

## Introduction

Embryogenesis, the transformation of a single zygote into a complex multicellular organism, is a highly orchestrated process of cell division, fate specification, and morphogenetic movement (Burton & Torres-Padilla, 2025). This process is guided by a robust gene regulatory network (GRN) that maps genotype to phenotype (Nance et al., 2005). For decades, developmental biologists have sought to unravel this map at single-cell resolution.

The nematode *Caenorhabditis elegans* serves as an unparalleled model system for this pursuit, primarily due to its completely mapped and virtually invariant cell lineage (Sulston & Horvitz, 1977; Brenner, 1974). This detailed lineage provides a ground-truth framework for studying cell fate decisions (Sulston et al., 1983). The advent of single-cell RNA sequencing (scRNA-seq) has provided the tools to measure the transcriptomic state of these individual cells, offering a path to deconstruct developmental trajectories (Tang et al., 2009).

However, despite the availability of foundational scRNA-seq atlases for the early *C. elegans* embryo (Tintori et al., 2016; Packer et al., 2019; Cole et al., 2024), a fundamental challenge persists: cell identity annotation ambiguity (Tintori et al., 2016). As cells progress through rapid divisions, the transcriptomic differences between closely related sister cells (e.g., ABala and ABalp) become minimal. Conventional annotation methods, which rely on hierarchical clustering and known marker genes, struggle to distinguish these cells (Tintori et al., 2016). This leads to the pervasive issue of merged labels, where distinct identities are collapsed into ambiguous groups (e.g., ‘ABalx’). Such ambiguity is a primary bottleneck, obscuring the precise gene expression programs driving distinct fate choices and preventing the construction of high-resolution developmental models (Cole et al., 2024).

We hypothesize that this ambiguity arises because conventional methods treat cells as independent observations, ignoring the crucial intercellular context in which they develop (Packer et al., 2019). Cell fate decisions are not made in isolation; they are heavily influenced by the cellular microenvironment through cell-cell interaction (CCI), mediated by ligand–receptor (L–R) signaling pathways like Notch and Wnt (Nance & Priess, 2002). For instance, Notch signaling between neighbor cells differentiates ABa vs. ABp fates at the 4-cell stage, and Wnt signaling influences the ABar division axis at the 8-cell stage. These observations suggest that integrating intercellular context could significantly improve annotation precision.

Therefore, we propose CHACAM, a computational framework that resolves cell identity ambiguity by integrating intrinsic gene expression with the extrinsic physical and molecular context of CCIs. We built a comprehensive curated *C. elegans* L–R database and combined it with a high-resolution 4D spatial atlas (Cao et al., 2020) to provide this context. CHACAM introduces a hierarchical attention mechanism and a novel “triangulation” strategy that leverages learned, contact-specific L–R signatures to disambiguate transcriptomically-similar cells. For example, to resolve an ambiguous “ABalx” cell, CHACAM can use the contact profile of a reference cell like MSa, which contacts ABalp but not ABala, to assign the cell identity based on the learned L–R interaction signatures. Cross-validation from multiple reference cells confirms that such triangulation yields consistent, robust identity calls. We apply CHACAM to published *C. elegans* embryo scRNA-seq atlases and demonstrate that it significantly raises the single-cell ratio, defined as the ratio of uniquely resolved single-cell identities to all annotated cell identities, in the early embryo stages by successfully resolving the previously merged cell identities.

Here, we will detail the methodology of CHACAM, from L–R database curation to the architecture of the model. We demonstrate its superior performance in resolving ambiguities in the Tintori et al. (2016) and Cole et al. (2024) atlases, leading to a higher single-cell ratio transcriptomic map of early embryogenesis. Moreover, CHACAM uncovers novel cell-specific marker genes and reveals biologically interpretable interaction patterns. The methodology is general and can be adapted to other single-cell datasets with spatial or interaction information, laying a foundation for more precise and mechanism-informed cell annotations in developmental and biomedical research.

## Methods

### CHACAM architecture

CHACAM is a computational framework that assigns cell identities by integrating each cell’s gene expression profile with neighbor-contact information (Figure 1). The model’s architecture employs hierarchical attention at two levels, a gene-level attention to quantify the transcriptomic similarity and a neighbor-level-contact-based triangulation to disambiguate the cell identities.

**Figure 1.**
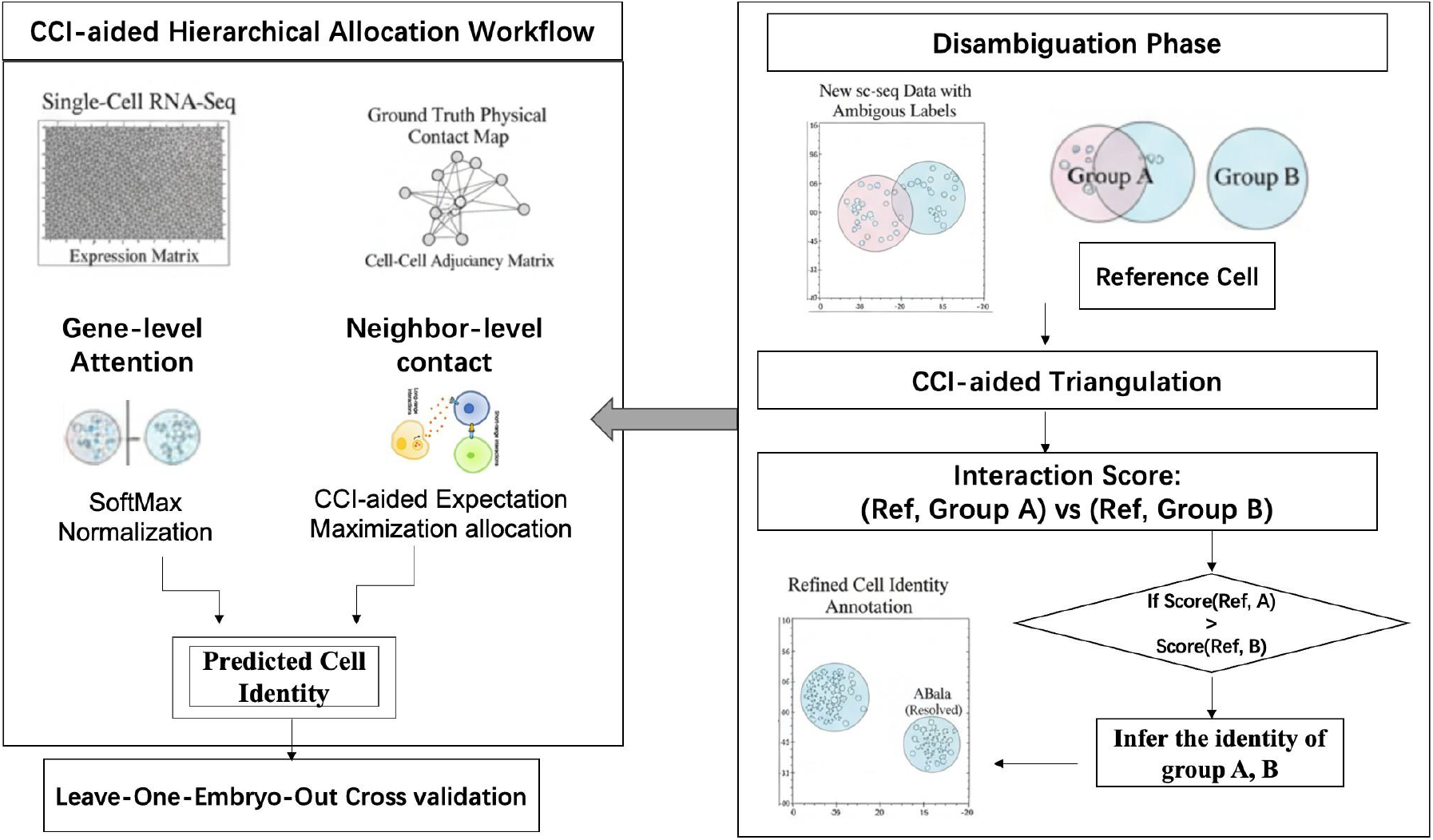
Overview of CHCAM workflow. Left: The pipeline integrates single-cell RNA-seq with a ground-truth physical contact (cell–cell adjacency) map. A gene-level attention module exploits discriminative transcriptomic features, while a neighbor-level module incorporates ligand–receptor interaction features from contacting neighbors. Outputs are combined to predict cell identities and are evaluated by leave-one-embryo-out cross-validation. Right: Disambiguation (triangulation) phase. A reference cell with known differential contacts is used to compare interaction scores [Ref-A vs Ref-B]. The score helps determine the identity, yielding a refined annotation.

First, a gene-level attention computes a cosine similarity of the cellular transcriptome between the target embryo and source embryo on the same set of key gene features (Figure 2). The similarity matrix is then normalized (e.g., row-wise and column-wise SoftMax) and combined (e.g., element-wise product) to produce the final attention score matrix *L*. The score *L*^ij^ represents the likelihood that target cell i corresponds to reference identity j. We next apply a greedy hierarchical allocation algorithm to assign a unique identity to each cell based on the attention scores, by locating the maximum attention score *L*^ij^ in the current matrix and assigning identity j to target cell i, then ignore the row i and column j and repeat until all cells are assigned.

**Figure 2.**
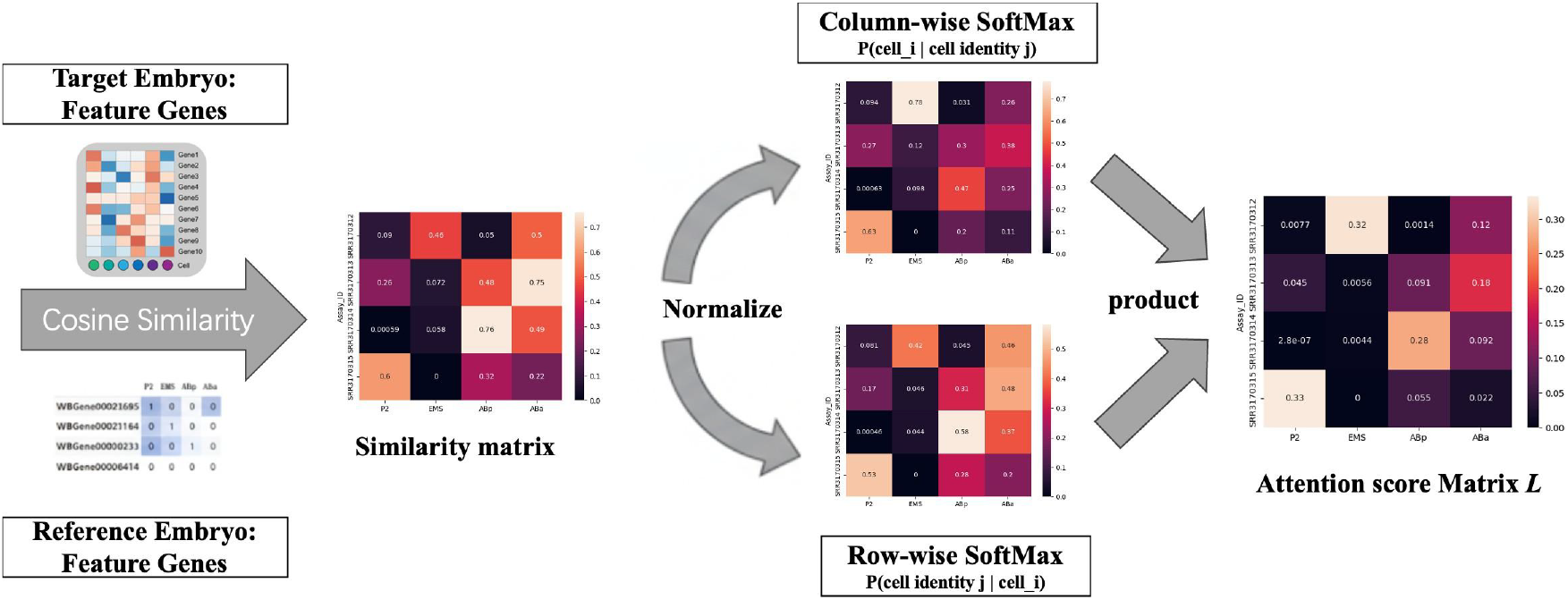
Schematic illustration of the attention score computation. The diagram illustrates the workflow for computing the expression-based attention scores, which forms the first stage of the CHACAM allocation model. **Inputs**: A Target Embryo (Query) and Reference Embryo (Key) are compared based on their expression profiles across a shared set of feature genes. **Similarity Matrix**: A matrix is generated by computing the cosine similarity between each target cell (Query) and each reference cell identity (Key). **Normalization**: The similarity matrix undergoes a bi-directional normalization. First, a Column-wise SoftMax is applied to calculate P(cell_i | identity_j). Second, a Row-wise SoftMax is applied to calculate P(identity_j | cell_i). **Final Score**: The two normalized probabilistic matrices are combined via an element-wise product to produce the final Attention Score Matrix L. This matrix provides a robust likelihood for matching a target cell to a reference identity, which is then used to guide the hierarchical allocation.

The standard hierarchical allocation could fail when the reference atlas itself is ambiguous (e.g., ‘ABalx’). For these cases, we developed a CCI-aided “triangulation” strategy by transforming the annotation from a simple similarity search into a constraint-satisfaction problem, where the assigned identity must be consistent with both the cell’s transcriptome and its physical interaction context.

### Triangulation for disambiguation

A key innovation in CHACAM is the triangulation strategy used to resolve ambiguously labeled cells, which uses a third cell with known contacts – a “reference cell” – to break symmetry between the two candidates.

we leverage known physical cellular contacts and choose a reference cell with confident annotation that has a differential interaction: it is known (from lineage and contact maps) to physically contact one of the two ambiguous identities but not the other. For example, to disambiguate ABalx, we can use the cell MSa as a reference, because MSa contacts ABalp but not ABala in the embryo (Figure 3). Using CHACAM’s learned interaction signatures, we can obtain the interaction score between the reference cell and each ambiguous cell (or each cluster of cells). The ambiguous cell (or cluster) that shows a higher interaction score with the reference is inferred to be the one that the reference does contact (e.g., ABalp), whereas the other is inferred as the non-contacted cell (e.g., ABala).

**Figure 3.**
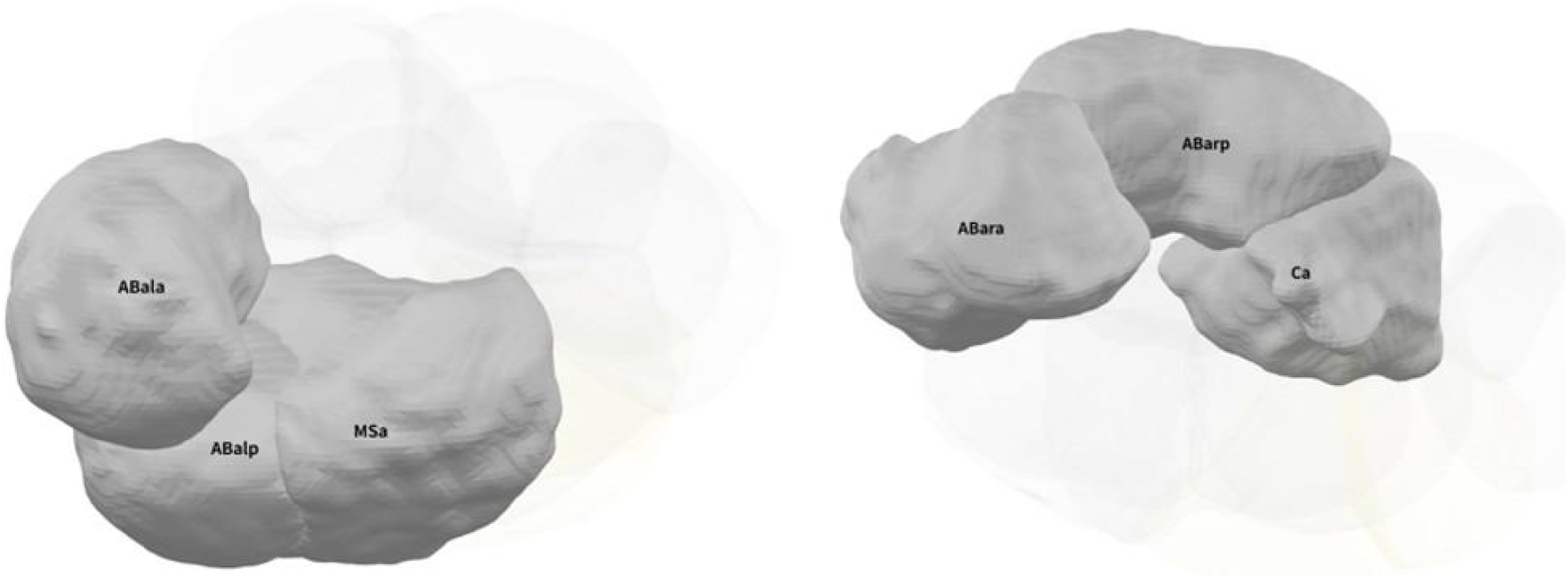
Example of triangulation strategy. Inferring Physical Cellular Contact between the pre-selected reference cell and the ambiguous ABalx & ABarx cells. MSa is selected as the reference cell to help distinguishing ABalx, because MSa is known to contact ABalp not ABala. Ca is selected as the reference cell to help distinguish ABarx, because Ca is known only to contact ABarp not ABara.

We applied this triangulation approach to each ambiguous pair when needed. In practice, multiple reference cells can be used sequentially or in combination to increase confidence. For instance, in addition to MSa for ABalx, one could also use MS if it had a known differential contact, and ensure both reference predictions agree. We evaluated the robustness of triangulation by cross-validation: using different reference cells and holding out embryos to verify that the assignments remain consistent.

## Results

### Improved Cell Lineage Resolution

#### Resolving ambiguous identities in early embryos

CHACAM achieved nearly perfect accuracy in annotating early embryonic cell identities, successfully resolving cases that were ambiguous in the original datasets.

In the Tintori C. elegans atlas (1-to 16-cell stages), our method assigned identity to every cell in leave-one-embryo-out evaluation, including those previously labeled with “x”. Notably, at the 16-cell stage, the 8 AB-lineage cells (ABala/p, ABara/p, ABpla/p, ABpra/p) had been originally collapsed into 4 indistinguishable pairs. CHACAM was able to fully disambiguate all of these: the single-cell resolution rate improved from 50% in the published annotations to 100%, restoring unique identities for all 16 cells at that stage (Figure 4). This constitutes the first time that a completely resolved lineage annotation (with no ambiguous cells) has been obtained for the 16-cell embryo in the Tintori dataset. All model-predicted identities were consistent with known lineage relationships and were confirmed by multi-reference cross-validation and held-out embryo tests.

**Figure 4.**
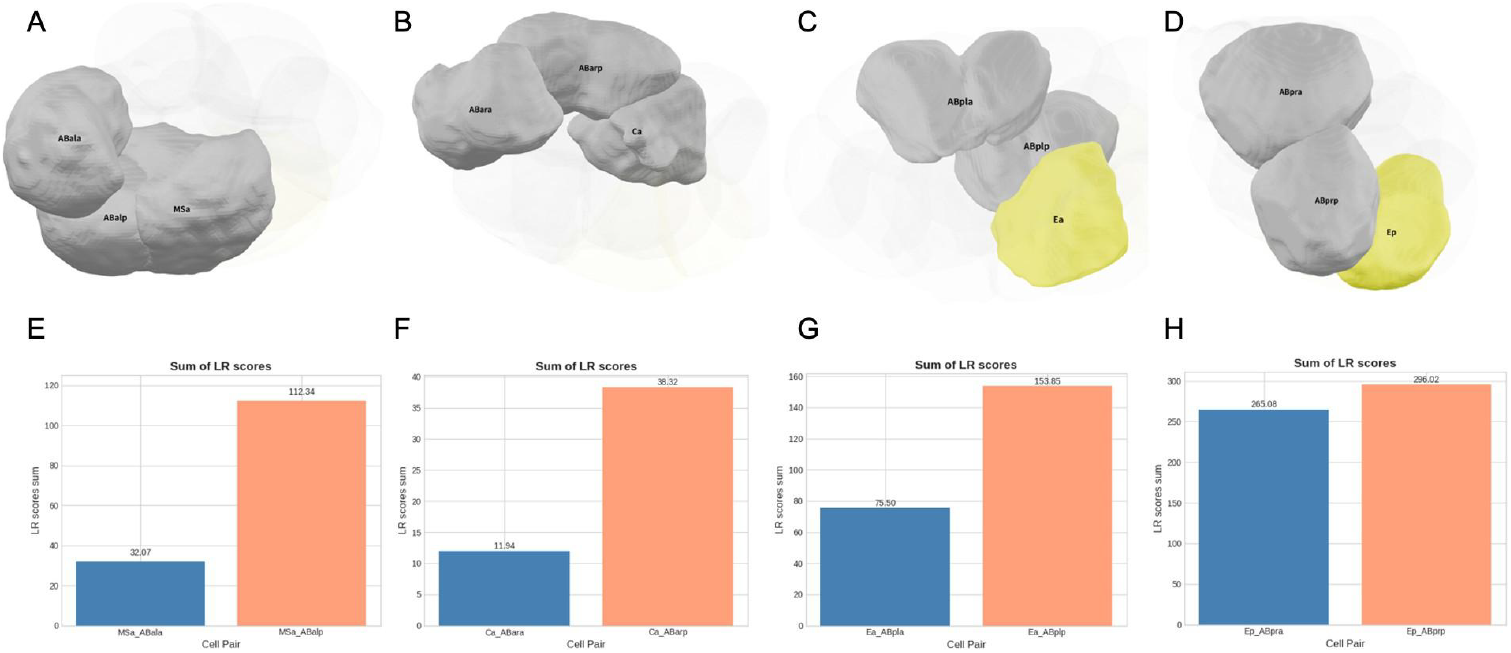
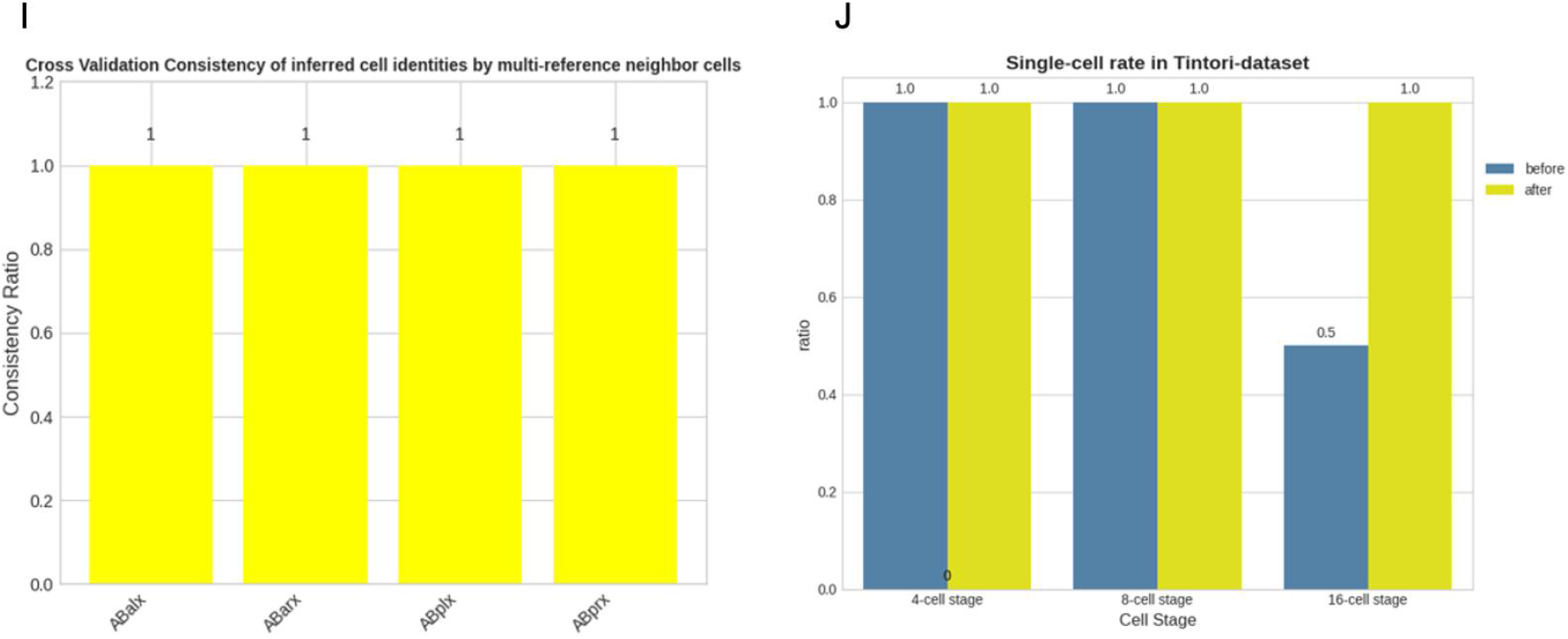
Results of CHACAM on Tintori-dataset. (A)-(D). Selected reference cells and the ambiguous ABxxx cells. (E)-(H). The total CCI score between the reference cell and ABxxx based on the top 50 contact-related ligand-receptors. The cell with higher score was inferred to be contact neighbor of the reference cell, the other to be non-contact. (I). The identity inference consistency across multi-reference cells. (J). The single-cell rate before / after annotations ambiguity resolved. It was increased to 100% at 16-cell stage in Tintori-dataset after ABxxx annotations ambiguity resolved.

In the Cole–Yanai dataset, CHACAM likewise maintained great annotation performance across the early and mid-stages. For embryos up to the 26-cell stage, where the original data had relatively complete sampling, CHACAM achieved essentially perfect single-cell resolution – all sister cells were distinguished. At later stages (e.g. 50-cell), the dataset itself contains some incomplete information (certain cells missing from the profiled data), which inherently limits achievable accuracy. Even so, CHACAM substantially outperformed an expression-only baseline, correctly annotating the vast majority of cells that were present. Overall, CHACAM’s accuracy was significantly higher than that of standard clustering or classifier approaches that ignore spatial context. These results establish that incorporating cell–cell interaction information leads to marked improvements in annotation precision, especially for challenging cases of sibling cells with similar transcriptional profiles.

### Discovery of Novel Marker Genes

Beyond improving annotation accuracy, CHACAM provided insights into gene markers that define each cell type. With all ambiguous cells resolved, we are now able to identify marker genes that discriminate each individual cell state. Many of these genes turned out to be novel marker candidates with cell-specific expression patterns. For example, in the 16-cell embryo, previous studies had used a handful of marker genes (such as tbx-38, hlh-27, ref-1and T09B4.1) to distinguish cells, but those markers could only partition the 8 AB-derived cells into 4 groups (Figure 5A). In contrast, the new identified a set of genes (such as gid-8, R144.10, lelo-4, and teg-1) exhibited unique expression in each of the eight AB blastomeres. Using these newly identified markers, all AB-lineage cells at the 16-cell stage can be individually recognized (Figure 5B).

**Figure 5.**
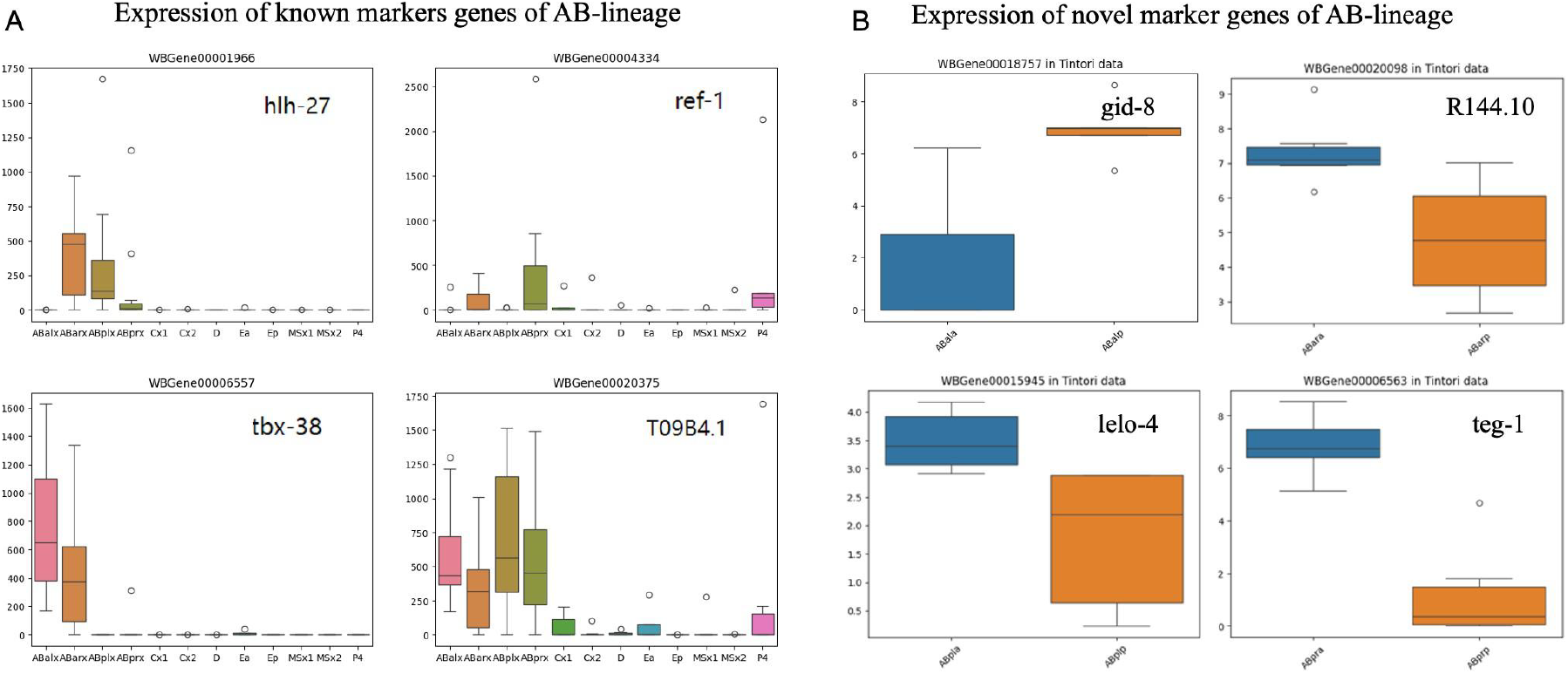
Prior-known and novel marker genes of ABxxx cells. (A). The expression of known marker genes (hlh-27, ref-1, tbx-38 and T09B4.1) used to annotate AB-lineage cells at 16-cell stage in Tintori-dataset. None of these genes are cell-specific, but the combination of them can partition the 8 AB-lineage cells into 4 groups: ABalx, ABarx, ABplx and ABprx. (B). The expression (log-transformed) of some novel marker genes that can further refine the AB-lineage cells into single-cell level. For instance, gid-8 can separate ABalp from ABala, R144.10 can separate ABara from ABarp, lelo-4 can separate ABpla from ABplp, and teg-1 can tell apart ABpra and ABprp.

We obtained similar results at other stages, many known lineage marker genes (e.g., pal-1 for the C/D lineages, med-2 for EMS) were indeed assigned high importance, validating the method. Importantly, however, the identified novel genes often have subtle yet distinct expression patterns that had been overlooked by manual analysis. For instance, certain chromatin factors and metabolic genes emerged as top discriminators for specific cells, suggesting previously unappreciated roles in those lineages. The full lists of marker candidates for each cell are provided in Supplementary files. These refined marker sets constitute a high-resolution transcriptomic atlas of the early embryo, which can serve as a resource for further hypothesis generation (e.g., testing the function of these genes in fate specification). In summary, CHACAM not only assigns cell identities with high confidence, but also helps explain those identities in terms of specific gene expression signatures.

## Discussion and Conclusion

Our results demonstrate that incorporating cell–cell interaction context is both feasible and highly effective for single-cell identification in developmental biology. CHACAM represents a shift from treating each cell in isolation to viewing each cell as part of a communicating community. In doing so, it resolves ambiguities that have persisted in scRNA-seq analyses of early embryos and delivers a more faithful molecular atlas of C. elegans embryogenesis. This work confirms that intercellular context can be decisive for accurate cell identity annotation. A cell’s identity is determined not only by its intrinsic gene expression program, but also by its extrinsic microenvironment – particularly the signals exchanged with its neighbors. By formally integrating this concept, CHACAM achieved a level of resolution that purely gene-based methods could not attain.

### Methodological implications

The success of CHACAM underscores the importance of bridging expression data with other data modalities (here, spatial contact maps and curated signaling knowledge). In developmental systems like C. elegans, decades of embryology have mapped out which cells touch or signal with which others. Our approach shows that such prior knowledge can be quantitatively combined with transcriptomic data to greatly enhance analysis. In essence, CHACAM leverages the invariant lineage and well-characterized signaling pathways of *C. elegans* as a scaffolding to improve predictions. The result is an annotated single-cell dataset that is consistent with both molecular data and embryonic anatomy. This high-fidelity atlas can now be used as ground truth for downstream computational modeling of embryogenesis (e.g., for building predictive models of cell fate or simulating embryonic development). More broadly, our work suggests that any time cells are difficult to distinguish by expression alone, incorporating orthogonal information (physical proximity, known interactions, lineage constraints) may be the key to disambiguation.

### Generalization to other systems

While we focused on C. elegans, the principles of CHACAM are applicable to other biological systems. Many single-cell datasets in development or disease suffer from analogous problems of closely related cell types that are hard to distinguish. For example, in mammalian organoid models or tissue development, progenitor cells often exist in niches where signaling from neighboring cells influences their fate. If spatial coordinates or contact probabilities can be obtained (through imaging or spatial transcriptomics), a CHACAM-like framework could integrate that information to improve cell type annotation. The triangulation strategy would be transferable to cases where a particular cell–cell interaction (say between a stromal cell and one subtype of tumor cell) could serve as a diagnostic feature to distinguish cell subtypes. As spatially-resolved single-cell datasets become more common, we envision context-aware annotation methods like CHACAM gaining importance. Our framework only requires a list of potential interactions and either a known or inferred adjacency map – resources that are increasingly available for tissues and organoids. Indeed, techniques like CellPhoneDB (Efremova et al., 2020) and CellChat (Jin et al., 2021) have accumulated L–R databases for human and mouse, and spatial transcriptomic technologies can provide contact estimates. Thus, one could apply the CHACAM-like approach to, for instance, resolve immune cell subpopulations in the tumor microenvironment by using knowledge of which cells physically interact in the tissue architecture. The demonstration in C. elegans serves as a proof-of-concept for this wider applicability.

### Limitations

Despite its advantages, CHACAM has certain limitations. First, its performance depends on the completeness and accuracy of the prior knowledge supplied. If the curated L–R database is missing critical signaling pairs, the model might not have the necessary features to tell those cells apart. We did literature curation to compile the C. elegans L–R list, but it is still possible that some developmental signals are not included or not yet discovered. Similarly, CHACAM assumes the physical contact map is reliable. In C. elegans, the contact map came from precise 4D imaging data (Cao et al., 2020), but in other cases contact information might be noisy or inferred. Errors in the contact adjacency (false positive or missed contacts) could lead to spurious or missed interaction features. In our study, we mitigated this by focusing on stable contacts and cross-validating results, but generally the quality of spatial data is a limiting factor. A related limitation is that CHACAM currently treats the interaction context in a static way – we did not incorporate dynamic time-series information or how interactions change over time. In principle, an extension of this model could include temporal dynamics (for example, using sequential models to track how a cell’s neighbors change as it moves or how signaling might induce delayed transcriptional changes). Our current approach is static and stage-specific, which is appropriate for early cleavage stages that are short and relatively time-invariant in terms of fate decisions, but future work could integrate temporal context for later stages.

Another practical limitation is the requirement of having some known contacts or reference interactions to perform triangulation. In C. elegans, it is straightforward to pick reference cells for each ambiguous case. In less well-mapped organisms, one might have to rely on probabilistic contact maps or statistically inferred spatial proximity (e.g. the approximated precision matrix or partial correlations). CHACAM’s algorithm could still be applied, but the certainty of reference-based identity calls might be lower. In such scenarios, one could incorporate multiple references or even treat the reference choice as an optimization problem (choosing the cell that maximizes difference in interaction score between two identity hypotheses). These are nuances that could be explored in follow-up studies.

Despite the above limitations, CHACAM provides a strong foundation for context-aware single-cell analysis. The method yielded a fully resolved lineage transcriptional atlas for the early C. elegans embryo – a resource that was previously unreachable using conventional methods. This atlas and the associated list of interaction features open new avenues of research. For example, one could now ask which specific L–R interactions are driving each cell fate decision, and design experiments to perturb those signals at precise times. The interpretable nature of CHACAM means it not only identifies cells, but also generates hypotheses about why a cell is a certain type (e.g., “Cell X is likely ABalp because it receives Notch from MSa and expresses gene Y”). These hypotheses can be tested by targeted perturbations in embryos, linking computational predictions to functional validation.

In conclusion, our study highlights the power of integrating molecular and physical data in single-cell analysis. CHACAM resolves long-standing annotation ambiguities in a hallmark developmental system, improving the fidelity of the data and enabling more accurate downstream modeling. As single-cell technologies move toward richer datasets(including spatial, temporal, and multimodal measurements), approaches like CHACAM will be instrumental in extracting coherent biological knowledge. By embracing both the “intrinsic” and “extrinsic” factors that define a cell, we can move closer to a holistic, mechanistic understanding of development and cellular identity.

## Materials

### Datasets

We analyzed two published scRNA-seq atlases of early *C. elegans* embryogenesis. The first is the Tintori *et al*. (2016) dataset, a full-length transcriptomic atlas from the 1-cell to 16-cell stages using Smart-seq2. The second is the Cole *et al*. (2024) dataset (Cole–Yanai atlas) covering developmental stages up to 102-cell using CEL-Seq2. Both datasets provide transcriptomes for each blastomere along with partial lineage annotations.

Notably, in their original annotations several cells were left ambiguous – labeled with an “x” in the annotation when transcriptional data alone could not distinguish the cells, like significant ambiguity in the 16-cell embryo AB-lineage of Tintori-dataset (e.g., ABalx, ABarx, ABplx, ABprx), it also exhibits such annotation ambiguity beginning after the 8-cell stage of Cole-Yanai dataset. These ambiguously labeled cells form the primary challenge that CHACAM aims to resolve.

To obtain ground-truth physical cell-cell contact information, we used the high-resolution 4D confocal imaging atlas from Cao et al. (2020). This atlas was generated using deep-learning-based 4D segmentation and provides precise 3D spatial coordinates and morphological shape for every cell at 1-minute intervals through the 350-cell stage.

### Ligand–receptor database curation

A robust CCI analysis requires a comprehensive, species-specific L–R database. We therefore extended a previously established C. elegans L–R database (Armingol et al., 2022) by incorporating additional interactions relevant to embryonic development. The previously curated L–R interactions were used as the foundation of our database, and additional candidate interactions were collected from primary literature and relevant resources, including WormBook, KEGG signaling pathways, and curated protein–protein interaction datasets. We specifically focused on interactions with known or potential roles in embryonic cell–cell communication, including key developmental signaling pathways such as Notch, Wnt, and TGF-β, as well as cell adhesion molecules.

After literature-based curation and removal of redundant interactions, the resulting database contained 286 unique L–R pairs, including 225 pairs retained from the previously established database and 61 additional pairs incorporated in this study. All gene and protein names were standardized to WormBase identifiers to ensure consistency in downstream analyses. This extended, embryogenesis-focused L–R database serves as the foundational interaction knowledge base for CHACAM, allowing the model to focus on biologically plausible interactions during early *C. elegans* development.

### Physical contact map

We integrated high-resolution spatial information by leveraging a 4D digital atlas of C. elegans embryogenesis (Cao et al., 2020). From these data, we derived a physical cell contact map for each developmental stage of interest. Two cells were defined as “in contact” if their cell surfaces were directly adjacent in the 3D embryo reconstruction at a given time point. We represented each stage’s contacts as a binary adjacency matrix, where entry (i, j) = 1 indicates that cell_i and cell_j are touching at that stage.

To focus on biologically meaningful contacts, we filtered out insignificant ones. We constructed a database of “stable” contact pairs by excluding contacts that occurred only for a short time (shorter than a specified duration, e.g., 2 minutes) or over a small surface area (less than a specified contact area ratio, e.g., 1/48). This ensured that our model emphasizes sustained cell–cell interactions rather than momentary juxtapositions due to cell random movement. The curated contact maps from the 4D atlas, in combination with the L–R database, form the basis for computing cell interaction features in CHACAM.

### Implementation and evaluation

We implemented CHACAM in Python using PyTorch for model building/training and Scanpy for data handling (Virshup et al., 2023) and LIANA+ for CCI analysis (Dimitrov et al., 2024). To evaluate performance, we conducted leave-one-embryo-out cross-validation on each dataset. That is, in each fold, data from one embryo (all cells across stages for that embryo) were held out as a test set, and the model was trained on the remaining embryos. This strategy assesses the model’s ability to generalize to new embryos (biological replicates) not seen during training.

Our primary evaluation metric was annotation accuracy: the fraction of cells in the test set whose predicted identity matches the true lineage identity. We computed accuracy at each developmental stage and overall. In addition, we examined the single-cell resolution rate, defined as the proportion of cells at a given stage that are uniquely identified (not merged). This metric is 100% if all cells at that stage receive distinct correct labels.

We compared CHACAM against baseline methods that do not incorporate cell–cell interactions. Its performance reflects what is achievable by transcriptional data alone. By contrasting this baseline with the full CHACAM, we quantify the added value of incorporating L–R interaction context. All models were trained with the same data splits and evaluated on the same test sets. Statistical significance of performance differences was assessed where applicable using paired tests across cross-validation folds.

## Acknowledgements

This work was partially supported by the Cecil H. and Ida Green endowment fund.

## Data Availability

The datasets analyzed in this study are publicly available from their original sources. The source code and curated data supporting this study will be made publicly available through GitHub.

## Author contributions

Xingyu Chen conceived the study, implemented the model and performed the analyses; Xinxin Ju and Maithri Murali helped with database curation; Michael Q. Zhang supervised the project and revised the manuscript.

## Supplementary Material

(Figures, database details, and marker gene tables placeholder.)

